# Timing between cortical slow oscillations and heart rate bursts during sleep predicts perceptual speed, but not offline consolidation

**DOI:** 10.1101/360867

**Authors:** Mohsen Naji, Giri P Krishnan, Elizabeth A McDevitt, Maxim Bazhenov, Sara C Mednick

**Affiliations:** Department of Medicine, University of California San Diego, La Jolla, CA 92093; Princeton Neuroscience Institute, Princeton University, Princeton, NJ 08540; Department of Cognitive Sciences, University of California Irvine, Irvine, CA 92697

## Abstract

Central and autonomic nervous system activity are coupled during sleep. Cortical slow oscillations (SOs, <1Hz) coincide with brief bursts in heart rate (HR), but the functional consequence of this coupling in cognition remains elusive. We measured SO-HR temporal coupling (i.e., the peak-to-peak interval between downstate of SO event and HR burst) during a daytime nap, and asked whether this SO-HR timing measure was associated with perceptual speed and learning on a texture discrimination task, by testing subjects before and after a nap. The coherence of SO-HR events during sleep strongly correlated with an individual’s perceptual speed in the morning and evening test sessions, but not with their change in performance after the nap (i.e., consolidation). We confirmed this result in two additional experimental visits, and also discovered that this association was visit-specific, indicating a reliable state (not trait) marker. Thus, we introduce a novel physiological index that may be a useful marker of state-dependent processing speed of an individual.

**Significance Statement:** Studies show that autonomic and central nervous system activity is coupled. For example, increases in heart rate follow cortical slow oscillations during sleep. However, the functional significance of this coupling for cognition is not understood. In three experimental visits, we show that the timing between these sleep events (the peak-to-peak delay between the slow oscillation and the heart rate burst) is highly correlated with waking perceptual processing speed. This reliable individual difference measure may be a useful marker of generalized processing speed.

## Introduction

As the brain shifts into deeper stages of non-rapid eye movement (NREM) sleep, neural firing becomes more synchronized, eliciting slow, cortical oscillatory rhythms that can be measured with scalp electroencephalogram (EEG). One of the predominant rhythms of NREM sleep is the slow oscillation (SO, <1Hz), which reflect underlying fluctuations between periods of neuronal activity (up states) and silence (down states) (Dang-Vu, et al., 2008). Autonomic nervous system (ANS) activity also changes as sleep deepens, undergoing a shift toward more parasympathetic dominance (as indexed by high frequency component of heart beat-to-beat intervals; 0.15-0.4 Hz), leading to progressive heart rate deceleration. Synchrony between SOs and cardiac autonomic activity during sleep has been previously reported (Jurysta, et al., 2003) (de Zambotti, et al., 2016), however the functional significance of these associations remains elusive.

Prior frequency-based analyses that averages across several minutes of sleep have reported that slow wave activity (0.5-4Hz) changes with and is preceded by parasympathetic activity (Jurysta, et al., 2003), (Brandenberger, Ehrhart, Piquard, & Simon, 2001). Additionally, seconds after spontaneous and evoked Stage 2 SOs (i.e., K-complexes), HR shows rapid accelerations followed by decelerations (de Zambotti, et al., 2016). On an even finer scale, heartbeat-evoked potentials (HEPs) have been measured during wake and sleep (Lechinger, Heib, Gruber, Schabus, & Klimesch, 2015), and waking HEPs predict visual detection (Park, Correia, Ducorps, & Tallon-Baudry, 2014). Further, the coupling between spontaneous pupillary fluctuations, which is controlled by sympathetic and parasympathetic activity, and brain activity during resting EEG was correlated with trait-level attention (Breeden, Siegle, Norr, Gordon, & Vaidya, 2017). More recently, increasing alignment of EEG-vigilance states and autonomic signals (HR and skin conductance) during resting state corresponded to stronger cortical inhibition (Ulke, et al., 2017). These findings suggest that 1) both the magnitude and timing between central and autonomic events are reliable indicators of CNS-ANS interaction, and 2) CNS-ANS interactions during sleep may be related to cognition. Here, we explore whether an individual’s processing speed may be reliably measured across modalities (waking behavior and sleep physiology) by examining the association between perceptual discrimination speed thresholds and the temporal coupling of EEG and electrocardiogram (ECG) events during sleep.

The goals of the current study were three-fold. First, we used high-resolution analysis of the relative timing of coupling of HR bursts and cortical SOs during sleep to obtain a marker of autonomic/central timing. Second, we sought to measure the association between two independent measures of processing speed: SO-HR timing and texture discrimination (i.e., the speed at which individuals can reliably discriminate the orientation of three target elements against a background of distractor elements), as well as the improvement in discrimination that is known to occur after a period of sleep (Mednick, Nakayama, & Stickgold, 2003). Third, we measured the reliability of the SO-HR and perceptual performance association across three experimental visits. Subjects were tested on three occasions, two weeks apart. On each experimental day, subjects completed a texture discrimination task (TDT) at 9AM and 5PM, and took a 90-minute polysomnographically-recorded nap (approximately 1:30PM to 3:30PM) between task sessions. The discrimination target was in a different visual field location at each visit, but constant within-xday. We found that SO-HR timing during the nap is associated with perceptual speed during each test session, but not necessarily with the change in performance across sessions. Furthermore, we found that this association is specific to the current state of the individual, as measured by poor cross-visit correlations.

## Materials & Methods

### A. Participants

Twenty-nine (16 females) healthy, non-smoking adults between the ages of 18 and 28 with no personal history of sleep disorders, neurological, psychological, or other chronic illness gave informed consent to participate in the study. All experimental procedures were approved by the Human Research Review Board at the University of California, Riverside and were in accordance with federal (NIH) guidelines and regulations. Participants included in the study had a regular sleep-wake schedule (reporting a habitual time in bed of about 7–9 h per night). Participants were thoroughly screened prior to participation in the study. The Epworth Sleepiness Scale (ESS) and the reduced Morningness-Eveningness questionnaire (rMEQ) were used to exclude potential participants with excessive daytime sleepiness (ESS scores >10) or extreme chronotypes (rMEQ <8 or >21). Participants received monetary compensation for participating in the study.

### B. Data acquisition and analysis

#### Study Procedure

Data reported here come from a larger, mini-longitudinal study that included up to 7 visits per participant. Data from this study have been reported elsewhere (Whitehurst, Cellini, McDevitt, Duggan, & Mednick, 2016) (Cellini, Whitehurst, McDevitt, & Mednick, 2016) (Sattari, et al., 2017). Subjects completed three in-lab study days, one each at the beginning (Visit 1), middle (Visit 2) and end (Visit 3) of the experimental period, spaced two weeks (14 +/- 2 days) apart. Participants wore actiwatches to monitor sleep-wake activity for one week prior to the experiment to ensure participants were not sleep-deprived and spent at least 6.5 hours in bed the night prior to their visit. Subjects arrived at the UC Riverside Sleep and Cognition lab at 9AM and completed the perceptual learning task (see below). At 1:30PM, subjects took a polysomnographically-recorded nap. They were given up to 2 hours time-in-bed to obtain up to 90 min total sleep time. Sleep was monitored online by a trained sleep technician. Nap sessions were ended if the participant spent more than 30 consecutive min awake. At 5PM, subjects were retested on the perceptual learning task. After excluding outliers based on behavioral performance (M ± 3SD), a total of 29, 28, and 25 subjects from Visit 1, Visit 2, and Visit 3, were used for the analyses, respectively.

#### Sleep recording

Polysomnography (PSG) recordings, which included electroencephalogram (EEG), electrocardiogram (ECG), chin electromyogram (EMG), and electrooculogram (EOG), were collected using Astro-Med Grass Heritage Model 15 amplifiers with Grass GAMMA software. Scalp EEG and EOG electrodes were referenced to unlinked contralateral mastoids (F3/A2, F4/A1, C3/A2, C4/A1, P3/A2, P4/A1, O1/A2, O2/A1, LOC/A2, ROC/A1) and two submental EMG electrodes were attached under the chin and referenced to each other. ECG was recorded by using a modified Lead II Einthoven configxuration. All data were digitized at 256 Hz.

#### Sleep scoring

Raw data were visually scored in 30-sec epochs according to Rechtshaffen and Kales (Rechtschaffen & Kales, 1968). Five stages (i.e., wake, sleep Stage 1, Stage 2, SWS, and REM) were reclassified in continuative and undisturbed 3-min bins, which were used for further analysis.

#### Heart-beat detection and time-series extraction

The ECG signals were filtered with a passband of 0.5-100 Hz by Butterworth filter. R waves were identified in the ECG using the Pan-Tompkins method (Pan & Tompkins, 1985), and confirmed with visual inspection. In order to extract continuous RR time-series, the RR intervals were resampled (at 4 Hz) and interpolated by piecewise cubic spline.

#### Slow oscillation

The EEG signals were filtered (zero-phase bandpass, .15–4 Hz). Then, SO were detected from the F3 and F4 electrodes based on a set of criteria for peak-to-peak amplitude, up-state amplitude, and duration of down- and up-states (Dang-Vu, et al., 2008).

#### -SO-HR Timing calculation

For each frontal electrode, an average RR time-series in reference to the down-state trough of SOs was calculated in a 10 s window. Then, the SO-HR timing was calculated by averaging the HR maximum times (i.e., RR minimum times) across the electrodes.

#### -Texture discrimination task (TDT)

Subjects performed a texture discrimination task similar to that developed by Karni & Sagi (Karni & Sagi, 1991). Visual stimuli for the TDT were created using the Psychophysics Toolbox (http://psychtoolbox.org). Each stimulus contained two targets: a central letter (‘T’ or ‘L’), and a peripheral line array (vertical or horizontal orientation) in one of four quadrants (lower left, lower right, upper left, or upper right) at 2.5°–5.9° eccentricity from the center of the screen. The quadrant was counterbalanced across subjects and visits. The peripheral array consisted of three diagonal bars that were either arranged in a horizontal or vertical array against a background of horizontally-oriented background distracters, which created a texture difference between the target and the background.

An experimental *trial* consisted of the following sequence of four screens: central fixation cross, target screen for 33 ms, blank screen for a duration between 0 and 600 ms (the inter-stimulus-interval, or ISI), mask for 17 ms, followed by the response time interval (2000 ms) and feedback (250 ms, red fixation cross with auditory beep for incorrect trials and green fixation cross for correct trials) before the next trial. Subjects discriminated two targets per trial by reporting both the letter at central fixation (‘T’ or ‘L’) and the orientation of the peripheral array of three diagonal lines (horizontal or vertical) by making two key presses. The central task controlled for eye movements.

Each *block* consisted of 25 trials, each with the same ISI. A threshold was determined from the performance across 13 blocks, with a progressively shorter ISI, starting with 600ms and ending with 0 ms. The specific sequence of ISIs across an entire session was (600, 500, 400, 300, 250, 200, 167, 150, 133, 100, 67, 33, 0). A psychometric function of percent correct for each block was fit with a Weibull function to determine the ISI at which performance yielded 80% accuracy. Within-day TDT performance change was calculated as the difference in threshold between Session 1 and Session 2, such that a positive score indicates performance improvement (i.e., decreased threshold in Session 2), whereas a negative score indicates deterioration.

Subjects were given task instructions and practiced the task during an orientation appointment prior to starting the study. During this practice, the peripheral target was located in a quadrant that was not used during the study. This practice ensured that subjects understood the task and accomplished the general task learning that typically occurs the first time a subject performs a task. Additionally, on the study day, subjects were allowed to practice an easy version of the task (ISI of 1000-600ms) prior to starting the test session to make sure subjects were able to discriminate the peripheral target between 90% and 100% correct on an easy version of the task.

## Results

For each visit, we used an automated algorithm to detect SOs for left and right frontal electrodes (F3 and F4, respectively) in Stage 2 and slow-wave sleep (SWS). We then analyzed heart rate activity within the ±5 second window following the SO downstate. During Stage 2 sleep, this window was characterized by an acceleration in heart rate following by a deceleration (Figure 1), in agreement with previous studies (de Zambotti, et al., 2016). The detected SO events during stable Stage 2 bins, co-occurred with a peak in heart rate, which was 12.09±1.48 % above the average Stage 2 heart rate. The average duration of the HR acceleration and deceleration was 5.52±2.51 sec. By comparison, during SWS, HR increased by only 3.35±1.01% following SOs.

**Figure 1.**
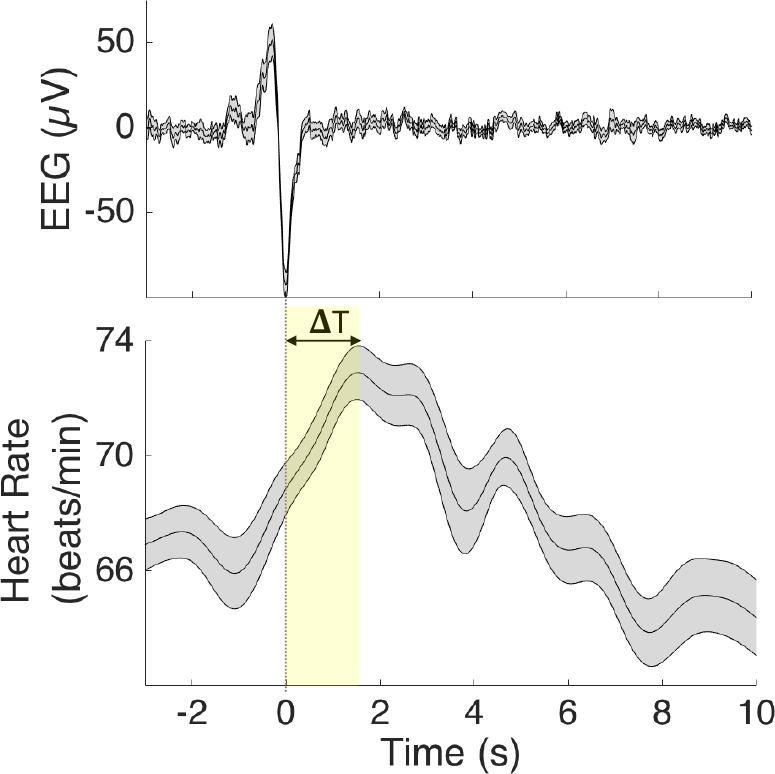
Heart rate reaches a peak following slow oscillations during Stage 2 sleep. The SO-HR timing is defined as the time difference between SO down-state trough and the heart rate peak.

We next measured duration of the SO-HR intervals. For each detected SO event in Stage 2 sleep, the SO-HR peak-to-peak interval was quantified by measuring the time difference between the trough of the SO down-state and peak of the average HR acceleration (Figure 1). The average SO-HR interval across F3 and F4 electrodes was used as the measure of SO-HR timing (ΔT). The average Stage 2 ΔT for visit 1, visit 2, and visit 3 naps was 2.15 s (s.d.=1.03), 2.10 s (s.d.=0.76), and 2.23 (s.d.=0.72), respectively. We next used each subject’s SO-HR timing at each visit to investigate its possible relation to perceptual speed as measured by TDT thresholds (pre- and post-nap sessions independently), and change in TDT performance after the sleep (change from pre- to post-nap sessions).

Pearson correlations revealed high correlation between pre-nap and post-nap performances within each visit (visit 1: r=0.89, p<.0001; visit 2: r=0.86, p<.0001; visit 3: r=0.80, p<.0001). Interestingly, SO-HR intervals were correlated with TDT performance in both pre- (visit 1: r=0.47, p=.01; visit 2: r=0.58, p=.001; visit 3: r=0.58, p=.002; Figure 2A) and post-nap sessions (visit 1: r=0.56, p=.002; visit 2: r=0.50, p=.007; visit 3: r=0.52, p=.008; Figure 2B) at each visit. On the other hand, there were no significant associations between SO-HR intervals and TDT performance change at any visit (visit 1: r=-0.07, p=.725; visit 2: r=0.175, p=.371; visit 3: r=-0.12, p=.561; Figure 2C). In order to confirm the visit-specificity of these associations, we investigated the correlations between the SO-HR timings in each visit and TDT performance in other visits. We found no significant cross-visit correlations (Figure 2D-E), suggesting that the relation between these two measures is specific to the current state of the individual, at least at the level of weeks. Lastly, we checked whether the SO-HR was simply a proxy for sleepiness by examining the correlation between SO-HR and subjective ratings of sleepiness. No significant correlation was found between the SO-HR timing and average sleepiness in pre-nap (visit 1: r=0.001, p=0.99; visit 2: r=0.01, p=0.97; visit 3: r=-0.06, p=0.78) and post-nap (visit 1: r=0.08, p=0.66; visit 2: r=0.06, p=0.74; visit 3: r=0.31, p=0.12) sessions. In summary, our data suggest evidence of a general marker of processing speed in the individual as measured by the association between SO-HR intervals during a nap and the speed of texture discrimination in the TDT. This finding was confirmed across all three experimental visits that were spaced two weeks apart. We did not find associations between our measure of autonomic/central coupling and sleep-dependent learning, and the cross modal association in processing speed was highly specific to the experimental

**Figure 2.**
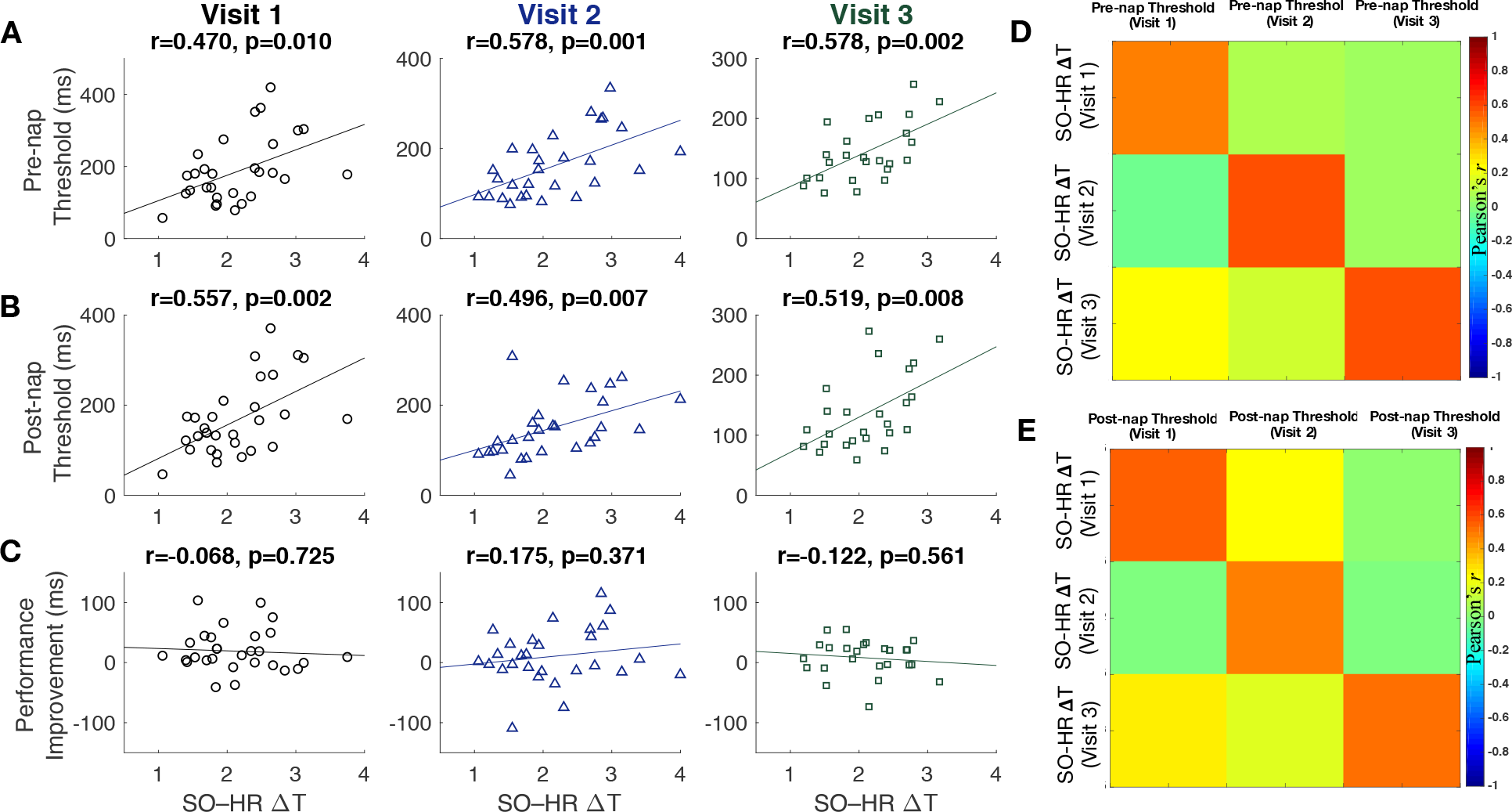
The scatter plots) show the relationship between SO-HR timing and A) Pre-nap TDT threshold, B) Post-nap threshold, and C) improvement in TDT performance. The lower threshold implies better performance. D-E) The cross-visit relationship between SO-HR timing and the TDT thresholds.

## Discussion

Prior work has shown a correlation between waking HEPs and attention (Park, Correia, Ducorps, & Tallon-Baudry, 2014). Furthermore, evidence has been shown for an association between cognition and other autonomic signals, including HR, skin conductance and gut-brain interactions (Tallon-Baudry, Campana, Park, & Babo-Rebelo, 2017). Here, we found that the timing between autonomic and central events during sleep gives a reliable individual difference measure that is associated with an independent measure of processing speed, namely texture discrimination. This is interesting for several reasons. First, the measure of perceptual performance and autonomic-central interactions were measured at different times and in different states of consciousness, which may suggest that this relationship reflects a more general marker of timing. Second, we found stronger within-visit than cross-visit associations between perception and heart/brain interaction, indicating a state-dependent measure of processing speed that is not simply a proxy for sleepiness. Third, it suggests that processing speed in other cognitive timing domains (i.e., motor, audition, tactile, etc.) may be predicted by ANS/CNS timing. Given findings of age-related decreases in cognitive processing speed (Kerchner, et al., 2012) and dampened autonomic nervous system activity (Stein, Barzilay, Chaves, Domitrovich, & Gottdiener, 2009), further studies should explore these associations in older adults, along with potential interventions to enhance cross-modal processing speed.

## Acknowledgements

This work was supported by grants from NIH (R01AG046646), ONR (MURI: N000141310672), and Young Investigator Prize to Sara C. Mednick.

